# Feasibility of cognitive neuroscience data collection during a speleological expedition

**DOI:** 10.1101/2023.12.12.571158

**Authors:** Anita Paas, Hugo R. Jourde, Arnaud Brignol, Marie-Anick Savard, Zseyvfin Eyqvelle, Samuel Bassetto, Giovanni Beltrame, Emily B.J. Coffey

**Author notes:** These authors contributed equally (Anita Paas, Hugo R. Jourde, Emily B.J. Coffey). (E.B.J. Coffey), www.coffeylab.ca (E.B.J. Coffey).

## Abstract

In human cognitive neuroscience and neuropsychology studies, laboratory-based research tasks have been important to establish principles of brain function and its relationship to behaviour; however, they differ greatly from real-life experiences. Several elements of real-life situations that impact human performance, such as stressors, are difficult or impossible to replicate in the laboratory. Expeditions offer unique possibilities for studying human cognition in complex environments that can transfer to other situations with similar features. For example, as caves share several of the physical and psychological challenges of safety-critical environments such as spaceflight, underground expeditions have been developed as an analogue for astronaut training purposes, suggesting that they might also be suitable for studying aspects of behaviour and cognition that cannot be fully examined under laboratory conditions. While a large range of topics and tools have been proposed for use in such environments, few have been evaluated in the field. We tested the feasibility of collecting human physiological, cognitive, and subjective experience data concerning brain state, sleep, cognitive workload, and fatigue, during a speleological expedition in a remote region. We document our approaches and challenges experienced, and provide recommendations and suggestions to aid future work. The data support the idea that cave expeditions are relevant naturalistic paradigms that offer unique possibilities for cognitive neuroscience to complement laboratory work and help improve human performance and safety in operational environments.

## 1. Introduction

Cognitive neuroscience and neuropsychology traditionally uses refined and highly-controlled laboratory-based research tasks (Nastase et al., 2020). This approach has been valuable to establish many principles of brain function and their relationship to behaviour, but task situations and the behaviour they evoke may differ greatly from real-life experiences. In both human and animal research, scientists are starting to explore how studying naturalistic behaviours can contribute to our understanding of brain function and behaviour (Shemesh and Chen, 2023; Smith, 2023). Of particular interest for studying complex human behaviour are situations that cannot be confined to a laboratory environment for practical, logistical, or sometimes ethical reasons. Such situations may take place in unique conditions, span many days or weeks, or even threaten participants’ safety (Mogilever et al., 2018; Stahn and Kühn, 2022). Understanding human behaviour and brain function under challenging conditions is broadly relevant to a range of safety-critical operations and professions (Manjunatha et al., 2020; Paas et al., 2022; Palinkas and Suedfeld, 2021; Paulus et al., 2009).

Speleology is a discipline concerned with the exploration, description, and study of caves and cave systems. It is generally carried out under conditions of isolation or disconnection from everyday life, exposure to unusual physical and sensory conditions, reliance on equipment, and technical challenges. Speleology imposes risks and stressors that increase fatigue, reduce cognitive capabilities, and can lead to mistakes, accidents, and injury (Mogilever et al., 2018). As such, speleological expeditions are an excellent training ground for people who work in extreme environments (e.g., astronauts; Sauro et al. (2021)), and could be useful to evaluate and optimize equipment interfaces and operational protocols, test the effectiveness of countermeasures to stressors, and conduct fundamental research in psychology, cognition, and neuroscience. Of particular relevance are circadian rhythm and sleep; interpersonal interactions; stress, decision-making, and risk-taking behaviour; and sensation and perception (Mogilever et al., 2018, Figure 1).

**Figure 1:**
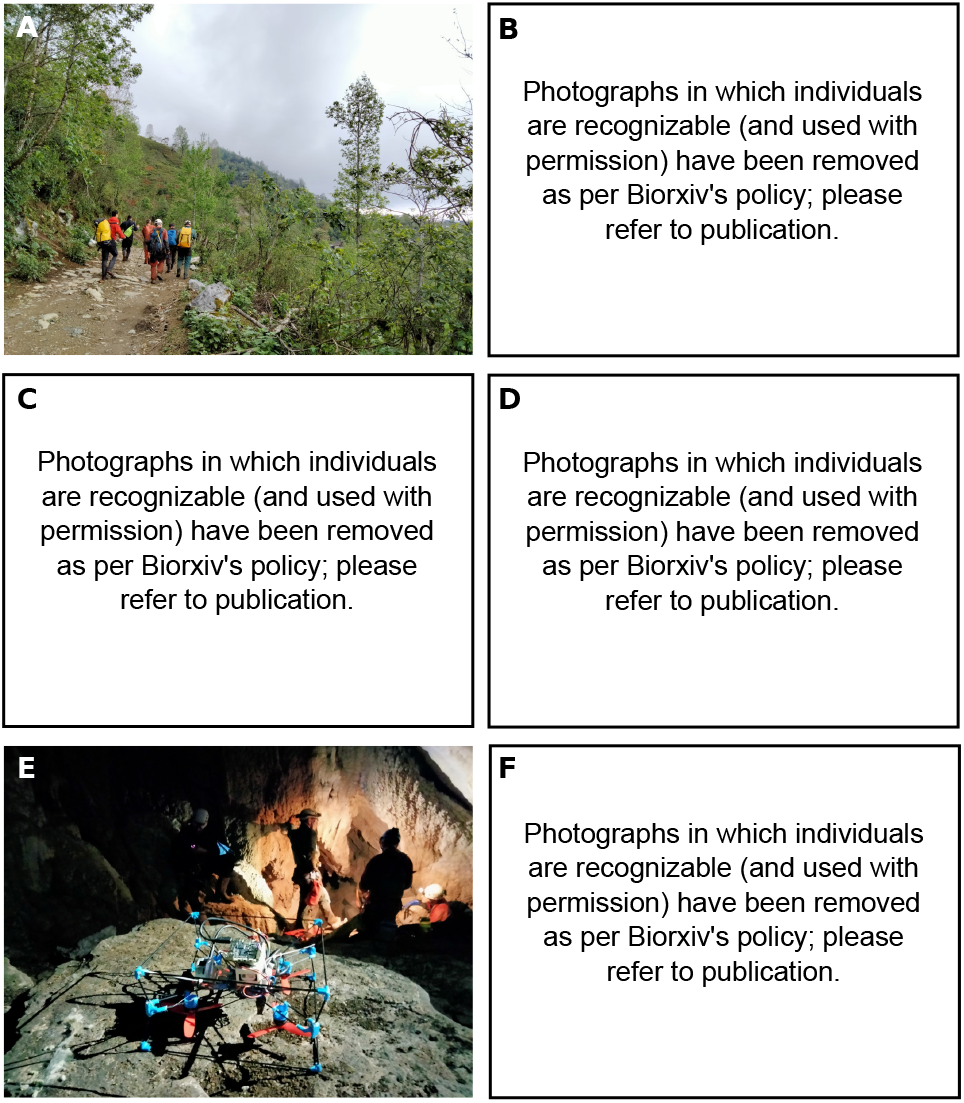
Conditions approaching and within caves. A) Caves were accessed on foot, in rugged terrain. Team members B) approaching a cave entrance, C) negotiating a vertical obstacle, and D) conducting mapping activities using commercially-available equipment. E) Two scientist-engineers tested a lightweight, collision-resistant drone, suitable for mapping the cave environment. F) A team member studies a 3D cave map produced using LIDAR scanning.

Conducting research on humans in extreme environments comes with its own set of challenges. For example, computerized tasks and other aspects of testing (questionnaires, interviews) must be planned and designed with consideration for the conditions, and the experimental setup needs to be portable and robust. Less experimental control means more variability in measured responses, and sample sizes are often small, generally meaning low signal-to-noise ratios of phenomena of interest. Tasks and stimuli must be chosen with an understanding of the environment and the other concurrent activities. For example, asking participants to complete time-consuming or effortful tasks while recovering from a full day of cave exploration may generate feelings of frustration or resentment, or even increase dropout rates (Mogilever et al., 2018). Furthermore, researchers must decide who will collect data, when, where, and how. There are many possible arrangements, such as: (a) scientists recording data remotely (e.g, by questionnaires or telephone-based interviews) during pre- and post-mission in-person meetings or in their own laboratories (e.g., examining changes in brain structure pre- vs. post-spaceflight; Jillings et al., 2020); (b) having participants gather data on themselves during an expedition (e.g., astronauts collecting data on sleep quality aboard the international space station; Koller et al., 2021); or (c) having the scientists on-site (e.g., observation of skydivers and climbers; Hardie-Bick and Bonner, 2016). The on-site approach (c) may be preferable when possible; several authors have emphasized that fieldwork benefits from first-hand experience and naturalistic observation of the phenomena under study (Douglas, 1976; Hardie-Bick and Bonner, 2016; Plate, 2007).

We tested the feasibility of embedded researchers collecting human physiological, cognitive, and subjective data during a speleological expedition in a remote region. We focused on fatigue and emotional regulation, as they are both related to cognitive performance and errors, and are therefore highly relevant for safety and mission success.

## 2. Methods

### 2.1. Expedition overview

Mexpé is a Quebec-based initiative that has taken place every few years since the 1980s. Each expedition involves ∼10-15 speleologists from Quebec (SpeléoQuébec), and sometimes other parts of Canada or abroad. Speleologists participate out of personal interest and are not remunerated. Expedition objectives relate to exploring and mapping an area in the south-eastern tip of the Mexican state of Puebla, around 300 km from Mexico City in the Sierra Negra mountain range, which contains previously unexplored caves. The exploration took place in rugged, hilly landscape (see Figure 1), in March 2023. In the specific region of exploration (∼>1000 m to 1800 m above sea level), temperatures were warm (18-26 C) and the climate was humid.

Conditions approaching and in the caves are illustrated in Figure 1. Caves are generally characterized by low light, high humidity, and cool temperatures. The caves varied in their degree of hydrological activity, airflow (which can produce a chilling effect in constricted spaces, particularly when clothing is wet), and types of obstacles (e.g., siphons, vertical shafts, subterranean lakes). During exploration activities, team members had no means of communication with the surface, base camp, or external world. The access to some of the cave entrances was sometimes physically-strenuous (∼500 m elevation gain on trails or in jungle, with equipment). Further description of expedition conditions is found in the Supplementary Material, and base camp conditions are illustrated in Figure 2.

**Figure 2:**
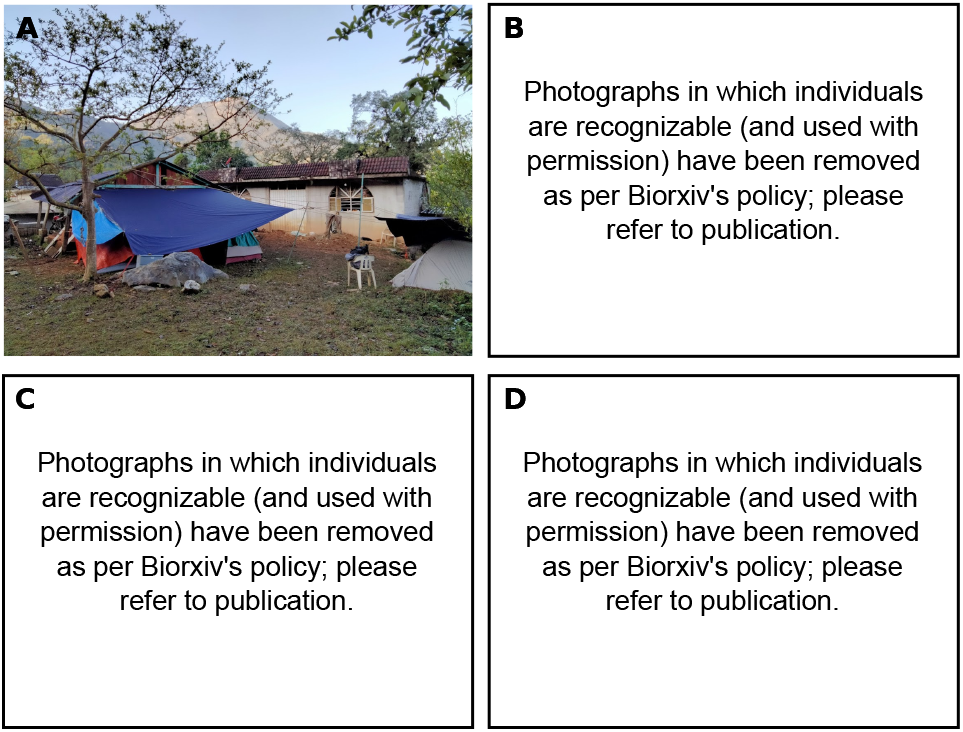
Base camp conditions. A) The camp comprised an abandoned building, an adjacent shed that was used for organizing and storing speleological equipment, and areas protected by tarps for tents. B) Heavy rainfall (and a lack of clean running water) challenged habitability of the camp environment. C) The main room, which was used for cognitive testing on Day 1, was also used intermittently for team meetings, conversation and meals, making it a distracting environment for data collection. D) On Days 2 and 3, cognitive testing was moved to a room reserved for sleeping, which provided a calmer and more predictable environment.

### 2.2. Participants

The group comprised 11 individuals, of whom 2 were female. Ages ranged from 27 to 65 years. Participants spoke a combination of English, French, and Spanish with different levels of fluency. All questionnaires, interviews, and tasks were conducted in English or French depending on each participant’s preferred language.

The majority of the speleologists had high to very high levels of speleological experience (i.e., most team members had been involved in speleology for 20-30 years, with one team member with 8 years and another 18 years experience). There were four embedded scientists and engineers; two focusing on cognitive neuroscience (i.e., current work) and two whose objectives involved testing three-dimensional mapping equipment. All four had general outdoor experience, three had some climbing experience, and one had previously been on a cave expedition as a course participant (similar to that described in Sauro et al., 2021). All four attended training on speleological equipment and procedures before the expedition, received refresher-training on-site, and were observed and guided by experienced team members while passing more difficult obstacles.

Embedded scientists were treated as participants, and non-scientist team members were encouraged to participate in the research process as collaborators; they were encouraged to make suggestions on the tools, relevance and feasibility of data collection. We note that using data collected on researchers may be inappropriate where knowledge of specific research hypotheses may influence the results (rather than exploratory questions). While unlikely here, we distinguish scientists from experienced cavers in plots, should differences arise that might be considered in future work.

Participation in both the expedition and in the scientific components was voluntary, and team members could opt out of data collection at any time. Written informed consent was obtained and study procedures were approved by the Concordia University’s Human Research Ethics Committee. Caving exploration permission and permits were obtained from local authorities to access the caves by expedition organizers, and goods and services were obtained on-site so as to support the local economy(e.g., a local shepherd with knowledge of the landscape was hired to guide the team to potential cave entrances).

### 2.3. Study design

The rationale for the selection of our measures is detailed in the Section 2 of the Supplementary Material. Figure 3 depicts the timeline of the scientific data collection, which took place during week 2 of the 3-week Mexpé 2023 expedition. Once on-site (Day 0), a team briefing ensured everyone understood the scientific objectives and procedures, and had an opportunity to discuss them with the researchers and leadership. Researchers prepared and tested equipment in situ and solved logistic issues (e.g., building a table for computerized testing, sharing access to three working electrical outlets). Cognitive and physiological data collection took place during the following three days, and sleep recordings were made on a subset of participants according to equipment availability. We elected not to study interpersonal interactions and social behaviour in the present work, and maintained a clear separation between data collection and ‘camp life’ periods, such that cave members did not feel under potentially unwelcome observation during recovery periods, and to better integrate the scientists within the team.

**Figure 3:**
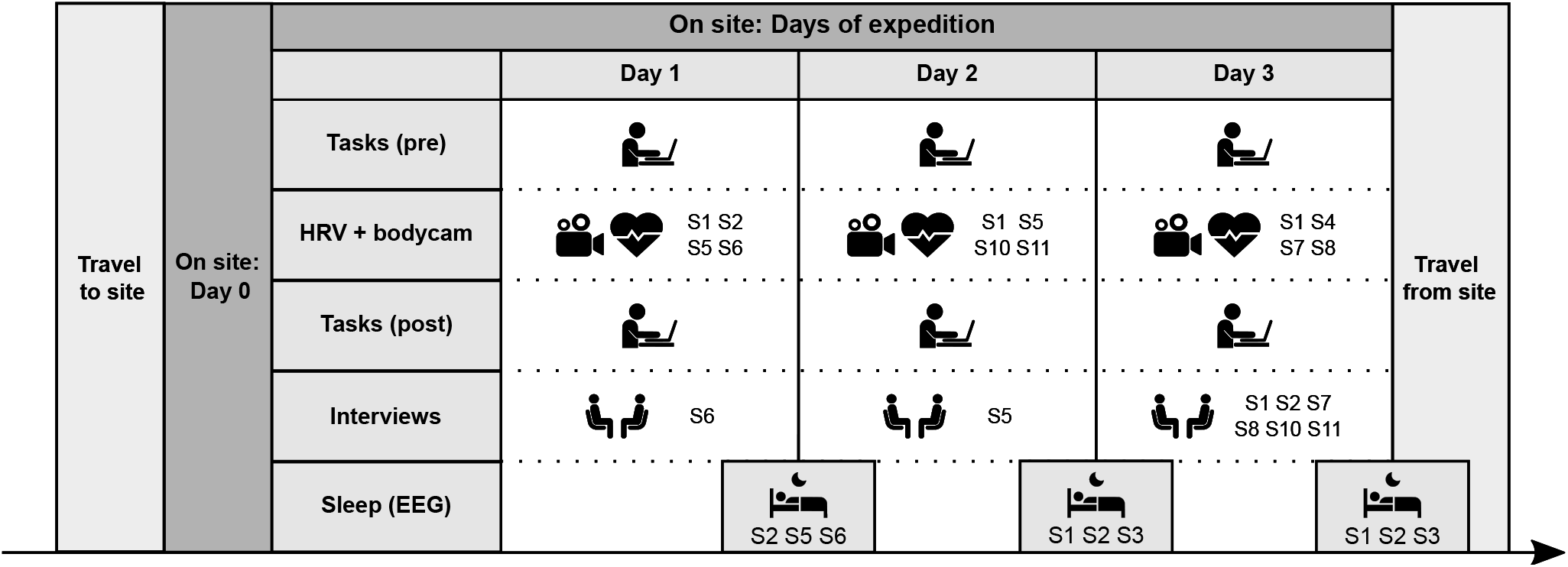
Overview of the four-day scientific portion of the expedition, starting with one day of organization, set-up, testing, and collection of questionnaire data (Day 0), three days of pre-post expedition cognitive testing and subjective fatigue ratings on all members and body camera and cardiovascular recordings on four team members at a time during exploration activities (Days 1-3). Sleep recordings were made on three participants per night (Days 1-3). One interview was conducted with each participant, mostly on the last day.

The cognitive testing setup is illustrated in Figure 2D. The team constructed a table from waste materials and placed laptops on it for cognitive testing. However, due to the very limited space (and rain on Day 1) and lack of furniture, the room was simultaneously used for meals, planning, and organizing equipment, making for a distracting environment (see Figure 2C). For Day 2 and 3, we moved the setup (see Figure 2D), and we used only these results in the analysis. Interviews during inclement weather were conducted in the storage room, and outdoors away from the main building during better weather. Daily fatigue ratings were obtained before or after cognitive testing when possible, otherwise the researchers tracked down team members during their morning and evening preparations to obtain ratings. Note that it was not always possible to obtain ratings without other team members being present.

The specific tests used and rationale for their adjustment for the expedition environment are discussed in the next sections, organized roughly according to mission phase: general questionnaires that were administered once on the first day (Day 0, see Single measurements); subjective and cognitive measures that were performed daily before and after exploration activities (see Daily measurements); physiological measures recorded within the caves (see Subterranean measurements); and sleep measures and interviews that were scattered throughout the expedition (see Occasional measurements). Due to space constraints, some detailed methodological descriptions are found in Section 3 in Supplementary Materials.

### 2.4. Single measurements

Questionnaires were answered on-site with pen-and-paper. As questionnaires were not previously available in French-language format, our bilingual team members translated them and reviewed the products for interpretation (translated versions used in the present work are available online^1^).

#### 2.4.1. Emotion Regulation Questionnaire

The Emotion Regulation Questionnaire (ERQ; Gross and John, 2003) is a 10-item scale designed to measure respondents’ tendencies to regulate their emotions with Cognitive Reappraisal and Expressive Suppression. Each item is answered on a 7-point Likert scale ranging from 1 (strongly disagree) to 7 (strongly agree). Items assess individuals’ emotional experiences (internal) and emotional expressions (external, such as gestures and behaviours). The Cognitive Reappraisal facet of emotion regulation is calculated by summing six of the items (for a total score between 6 and 42) and the Emotion Suppression facet score consists of the remaining four items (for a total score between 4 and 28).

#### 2.4.2. Psychological Skills Inventory for Caving

To document and explore the psychological skills of cave participants, we adapted a sports questionnaire (the Psychological Skills Inventory for Sports; Wheaton, 1998) on the basis that it might assess factors common to both activities, such as risk profiles, decision-making abilities, preparedness, and general skills and capabilities. The original scale comprised 60 items answered on a 5-point Likert from 0 (never) to 4 (always), and assessed six sets of skills: achievement motivation, goal setting, anxiety control, maintaining confidence, concentration, and mental rehearsal. Answers for each subscale are summed, and scores are calculated as a percentage of the total possible score for the subscale. Of the original items in the questionnaire, nine were removed because they could not be reformulated in a way that could refer to caving expeditions (e.g. ‘If I get behind in a competition, I feel that winning is impossible’). The rest of the items were reformulated so that any mention of “sport” and “competition” was changed to “caving” and “caving expedition” (e.g., ‘I set specific goals for every competition’ to ‘I set specific goals for every caving expedition’). On some items, the wording was simply changed to be more appropriate for a caving setting (e.g., ‘I am able to bounce back quickly after a disappointing performance’ to ‘I am able to bounce back quickly after making a mistake’). The reduced questionnaire had 51 items (7 related to ‘achievement motivation’, 6 for ‘goal setting’, 8 for ‘maintaining confidence’, and 10 items for each other subscale). As the modified questionnaire had not been used with speleologists, our testing focused on the relevance of questions to the speleologists. We invited participants to note questions they had difficulty answering, or found problematic or irrelevant, so as to generate an updated version for future work (provided in Supplementary Materials).

### 2.5. Daily measurements

#### 2.5.1. Subjective Fatigue Scale

Participants’ fatigue level was documented before and after exploration activities using a simple seven-item questionnaire that has been used in previous work (the Subjective Fatigue Scale, or SFS; de la Chapelle et al., 2022). The original questionnaire comprised 7 questions asking about different aspects of mental and physical fatigue rated from 0 (not at all) to 5 (very). We averaged the responses to 5 questions (tiredness, feeling active, feeling well, motivation, concentration; reverse-scoring the positively worded questions such that higher scores indicate greater levels of fatigue) to create a more global, composite measurement of general fatigue level. We omitted the questions ‘How relaxed are you?’ and ‘How interested are you?’ as some participants seemed confused by them or answered identically each time. These data were used to quantify changes in perceived fatigue before and after daily activity, and to compare with objective measures of cognitive performance.

#### 5.2.2. Cognitive tests

Both tasks ran in PsychoPy v2023.1.2. Tasks were performed in the morning between breakfast and departure (between 8 and 10 AM), and in the evening after participants had removed their climbing gear and taken care of basic needs (between 7 and 11 PM). Each task session lasted between 5 and 10 minutes, with the order of tasks counter-balanced. Participants were tested sequentially in groups of two.

##### Go/No-Go Task

In the Go/No-Go task, participants must press a key (the space bar) in response to a visual stimulus associated with the ‘go’ instruction (blue circle with the word ‘GO’ on it), and inhibit their response when presented with a less frequent visual stimulus associated with the ‘NO GO’ instruction (orange circle with the words ‘NO GO’ on it). The task was adapted from a version available online^2^. We changed the background from grey to black, to increase visibility of the stimuli through a higher stimulus-background contrast. The task comprised 100 trials (80 ‘GO’, 20 ‘NO GO’), each 1000 ms in length. The task therefore took less than two minutes. The visual stimulus was presented only during the first 750 ms of each trial. From this task, we obtained measures of error rate on the ‘NO GO’ trials (omission of the response) and average reaction time on the ‘GO’ trials, and subtracted participant means to facilitate comparison of differences across measurements. Participants were instructed to go as quickly as they could, but to prioritize accuracy.

##### Simon Switching Task

In the Simon Switching Task, participants must respond to a visual stimulus (arrow) presented on the computer screen, by pressing either the ‘z’ (left) or ‘m’ (right) key on the keyboard, according to two sets of instructions (depending on the colour of the arrow). If the arrow was orange, participants had to indicate whether the arrow was located on the left or right side of the screen, regardless of which direction the arrow was pointing. If the arrow was blue, participants had to respond based on the direction of the arrow, regardless of its location. The task measures the cost in performance that occurs during incongruent trials (i.e., when the location and direction of the arrow do not match), which is referred to as the “Simon Effect”; and the cost in performance for trials that occur after a change in instructions (i.e., switching from instructions for orange arrow to blue or vice versa), referred to as the “Switch Effect” (Simon, 1990). We subtracted the means of reaction time and accuracy of the congruent and non-switching trials from the means of the incongruent and switching trials, respectively, to produce the Simon and switch effects on accuracy and reaction time.

The task was adapted from a version available online^3^ . The colours of the arrows were changed to blue and orange (originally red and green) to accommodate possible red-green colour-blindness. The number of trials was increased from 48 to 100, with an equal number of trials in each condition. Trials lasted until the participant pressed one of the response keys, or after 4000 ms. The task took about three minutes to complete. Participants had access to a written reminder placed near the computer, in case they forgot the association between colour and instruction, and keys were labelled. They were instructed to go as quickly as they could, but to prioritize accuracy. Scores were normalized across subjects for comparison with subjective fatigue ratings, as in the Go/No-Go task. The Simon and switch effects are expressed in percent accuracy, and seconds for reaction time.

### 2.6. Subterranean measurements

The main goal of this phase of the data collection was to evaluate tools that could be used during cave exploration to assess cognitive workload, and in the future, potentially other topics such as errors, communication, and group interaction. We combined three measures: audiovisual recording on a body-mounted camera to later mark time periods of interest for analysis, heart rate monitors to record cardiac function (time-synchronized to the audiovisual recordings), and periodic subjective ratings. In which several participants were asked to report ratings of workload, arousal and valence (WAV) on a scale of 1 to 5 orally, and by holding fingers in front of the camera to represent their ratings.

These measures were selected in consultation with speleologists to ensure safety and minimal intrusion to ongoing activities. The camera and heart rate monitor were distributed to team members from different exploration groups, who had a range of tasks; for example, exploring the forest for cave entrances, removing equipment from previous exploration days, exploring new caves, mapping, or testing drone-based and hand-held mapping equipment and software. The scientists instructed participants on the use of the equipment in the morning prior to daily activities and demonstrated how to start the recordings.

### 2.7. Occasional measurements

#### 2.7.1. Sleep

We used three Dreem^4^ headbands to monitor neurophysiological correlates of sleep quality in a subset of participants during the expedition. One participant (S2) used the same headband for three consecutive nights (as well as three nights prior to the expedition for comparison), while the other two headbands were distributed to different participants (see Figure 3).

We qualitatively evaluated EEG signal quality, and compared subjective reports of sleep and environmental conditions with the recordings obtained using the headbands.

#### 2.7.2. Interviews

We adopted a semi-structured qualitative interview approach to capture participants’ experiences concerning their background, motivation for engaging in speleological activities, and how they think, feel, and act in extreme environments. Questions were developed in consultation with a researcher knowledgeable in clinical interview approaches (available online). The interview was divided into three parts. In the first, conducted once, participants were asked about their background and caving experience. The second part was intended to record observations about each days’ exploration activities, but due to time constraints was only conducted once per participant. The final part intended to look through the days’ body camera recordings to identify time points of interest, and asking participants to rate how they felt at that time (according to the WAV rating system). Interviews were recorded on the body cameras in audio-only mode. Following the expedition, interviews were transcribed for qualitative analysis.

## 3. Results

The results presented in the following sections are focused on addressing questions about feasibility, and developing recommendations for improving the measures or their administration for similar expedition environments, which are summarized in Table 1. The small sample size limits the generalization of the findings, but trends are noted for considering potential follow-up studies.

**Table 1.** Summary of scientific measures use in the Mexpé 2023 expedition, suggested applications, and recommendations for use.

| Measure | Purpose and application | Considerations |
| --- | --- | --- |
| ERQ | Document tendencies to use emotional regulation strategies | Easy to administer<br>Difficult to compare a single overall/trait across people<br>Questions on less relevant emotions (e.g. sadness) |
| PSIC | Document cave-related psychological skills | Easy to administer<br>Updated questions for cavers<br>Needs validation in larger sample |
| Subjective fatigue scale | Assess subjective fatigue state | Fast for repeated administration<br>Appears to relate to cognitive task results<br>Needs validation |
| Go/No-Go task | Measure changes in inhibition and speed-accuracy trade-offs | Feasible to repeat if kept short (2-3 minutes)<br>Appears to relate to subjective fatigue levels<br>Requires a practice session and clear instructions on accuracy-speed trade-offs |
| Simon Switching task | Measure changes in cognitive flexibility | Feasible to repeat if kept short<br>Appears to relate to subjective fatigue levels<br>Similar to Go/No-Go but focused on different cognitive aspects |
| A/V recordings | Marking periods of interest for physiological analysis<br>Contextualizing measures<br>Optional review a posteriori<br>Optional reuse of excerpts as stimuli for in-lab experiments | Infrared recording and audio of good quality<br>Needs stabilization<br>Can be difficult to interpret a posteriori, facilitated by logs<br>Requires trust and ethical treatment of data<br>Useful to have participants vocalize what they are seeing |
| Physiological measures | Objective measure of physiological state | HRV metrics can distinguish cognitive state and show trends<br>Can be confounded with activity levels<br>Is an established measure (sport watches, bands...)<br>Needs synchronization with video |
| WAV | Subjective measure of psychological state during exploration<br>Comparison with physiological measures | Effective and minimally intrusive<br>Needs an alarm to notify the user to provide regular ratings |
| EEG / abbreviated PSG | Objective measure of sleep physiology<br>Related to fatigue and cognitive performance | Works even in difficult conditions<br>Useful to document the severity of sleep disturbance and for circadian rhythm studies<br>Needs charging and data upload via internet connection<br>Basic research questions could be answered with smart watch data |
| Semi-structured interview | General background of individuals<br>Individuals strengths or challenges<br>Contextualize daily activities<br>Identify interests and concerns of speleologist collaborators | Daily interviews and video analysis too time-consuming on-site<br>Background experience can be done in advance<br>Can reveal important experiences and phenomena for future study<br>More revealing insights after trusting relationship is established |

### 3.1. Single measurements

#### 3.1.1. Emotion Regulation Questionnaire

The scores on the cognitive reappraisal subscale ranged from 21 to 41, with a mean of 30.7 (SD = 6.05, see Figure 4A). These scores seem somewhat different from those of community samples, which tend to show lower means in larger samples (between 28 and 29 with SD between 6 and 7 Melka et al., 2011; Preece et al., 2021, 2019). The scores on the expressive suppression subscale ranged from 5 to 23, with a mean of 13.41 (SD = 5.65, see Figure 4B), suggesting a trend towards lower means than in larger community samples (e.g., >15 with SD >6 in Preece et al., 2019). In our sample, the embedded scientists showed similar results to the cavers, except on the expressive suppression subscale, in which they seemed to score generally lower (Mann Whitney U test: *U* = 24.00, *p* = .072). We confirmed that the two subscales were not significantly correlated (*r* = −0.017, *p* = .96), in line with the theory that these constructs are orthogonal and do not overlap (Gross and John, 2003).

**Figure 4:**
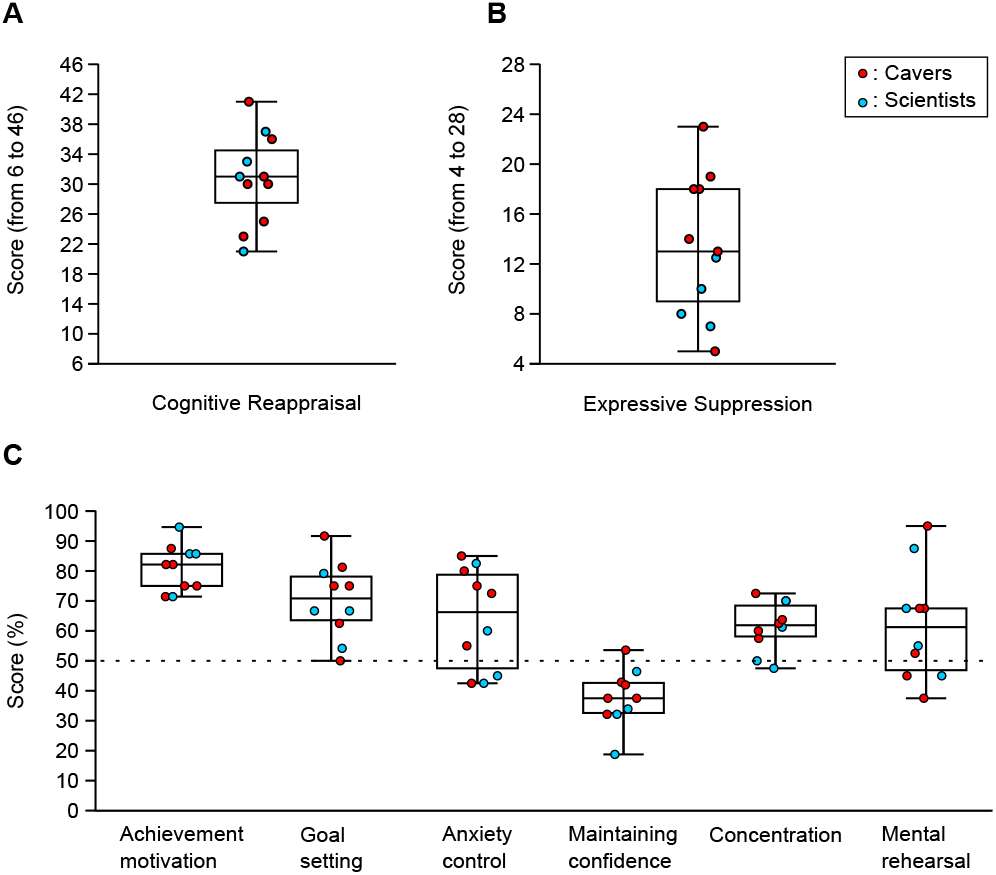
Team results on the Emotional Response Questionnaire (N=11): A) cognitive reappraisal and B) expressive suppression subscales, and C) on the Psychological Skills Inventory for Caves (N=10), a questionnaire we adapted from sports applications for use with cavers. The embedded scientists showed similar results to the cavers, except on the expressive suppression subscale, in which they scored generally lower. Of the six original subscales, the ‘Maintaining Confidence’ construct seems least applicable to caving expedition (almost all scores below 50%).

#### 3.1.2. Psychological Skills Inventory for Caves

The results of the PSIC should be interpreted with caution due to its status as a newly adapted questionnaire for the caving context, and because modifying and adapting the questionnaire introduced changes to the original scale. In the paper version of the questionnaire, the ‘sometimes’ column of the 5-point Likert scale was inadvertently omitted. Participants either answered from the existing columns (possible scores: 0, 1, 3, or 4), or marked a middle ground between two columns. Given that the potential maximal score for each subscale remained unchanged (4 x number of items), we could still derive scores (in percentage) by summing the item scores and dividing by the maximal score of each given subscale. We were able to derive some approximate scores for all subscales of the PSIC, allowing us to examine relative differences and patterns within and across subscales. One participant completed only half of the questionnaire, and was therefore excluded from figures and analyses.

Figure 4C shows the distribution of scores on each subscale, in addition to their respective median and quartiles. Expedition members scored higher on the achievement motivation subscale (M = 81.07, SD = 7.68) than all other scales, followed by goal setting (M = 70.83, SD = 12.73), anxiety control (M = 66.25, SD = 17.05), concentration (M = 61.88, SD = 8.29), and mental rehearsal (M = 61.25, SD = 18.70). Notably, the highest variability was in mental rehearsal, which was also highlighted in the feedback from participants (specifically, some participants noted that the ‘mental rehearsal’ items did not apply to them, while others had no issues with the questions). Finally, the subscale related to maintaining confidence showed the lowest scores, all below 54%, and seemed the least applicable to caving (M = 37.50, SD = 9.45).

These findings provide preliminary insights into the psychological skills of cave participants but warrant further validation and refinement to ensure the questionnaire’s robustness and suitability for future use in the speleological setting.

### 3.2. Daily measurements

#### 3.2.1. Subjective Fatigue Scale

Individuals’ subjective ratings of fatigue across the three experimental days are presented in Supplementary Figure 1. Unsurprisingly, participants reported greater levels of fatigue at the end of the day, collapsed across Days 1-3 (mean morning: 2.2, SD: 0.6; mean evening: 2.6, SD: 0.7; t(32) = 2.34, *p* = .026 [two-tailed]; Cohen’s d: 0.41). Participants reported moderate levels of fatigue in the mornings (Day 1 mean: 2.0, SD: 0.6; Day 2 mean: 2.0, SD: 0.5; Day 3 mean: 2.5, SD: 0.6; note that the scale range is from 1 to 5), likely due to challenging sleeping conditions. A repeated-measures ANOVA did not reveal a significant change in fatigue level across the three experimental days (F(2, 20) = 1.60, *p* = .2, *η*^2^ = 0.14). Knowledge of the teams’ roles, experience level and specific daily activities was important to interpret specific scores. For example, the scientific team, who were less experienced, generally showed greater increases in fatigue levels; a person who felt unwell on Day 1 stayed at the base camp to rest and reported lower fatigue levels in the evening.

#### 3.2.2. Cognitive tasks

Overall accuracy and general reaction time for both cognitive tasks for Days 2 and 3 are presented in Table 2 (one trial for one individual was removed as an outlier, >1.5 inter-quartile rule). We used Friedman tests (non-parametric repeated measures ANOVA) to analyze daily and pre/post expedition differences. We found a significant difference between daily pre- and post-reaction time values on the Simon Switching task (*χ*^2^ = 19.26, Kendall’s w = 0.35, *p* = .002), with post-hoc analyses showing differences in reaction time between first day pre-expedition and all other testing sessions (*p* < .05 for all tests). No such trend was observed for accuracy (*χ*^2^ = 6.61, w = 0.13, *p* = .25). Significant differences were not found in reaction time (*χ*^2^ = 3.21, w = 0.06, *p* = .67) or error rate (*χ*^2^ = 2.38, w = 0.04, *p* = .79) for the Go/No-Go task, possibly reflecting its simpler nature with fewer required executive functions.

**Table 2.** Go/No-Go and Simon Switching Tasks Descriptive Statistics.

|  | Go/No-Go task |  | Simon Switching task |  |
| --- | --- | --- | --- | --- |
|  | Error rate (%) | Reaction time (s) | Accuracy (%) | Reaction time (s) |
| Mean | 13.86 | 0.36 | 96.25 | 0.76 |
| St. Dev. | 14.59 | 0.05 | 3.42 | 0.24 |
| Minimum | 0 | 0.27 | 84 | 0.44 |
| Maximum | 50 | 0.49 | 100 | 1.45 |

Greater fatigue levels seemed to decrease reaction time the Go/No-Go ask (Figure 6B), with more modest in-eases in error rate (Figure 6A). Taken together with the observed relationship between error rate and reaction time (Figure 5), the results suggest that fatigue levels can affect people’s trade-off between accomplishing tasks quickly or with as few errors as possible.

**Figure 5:**
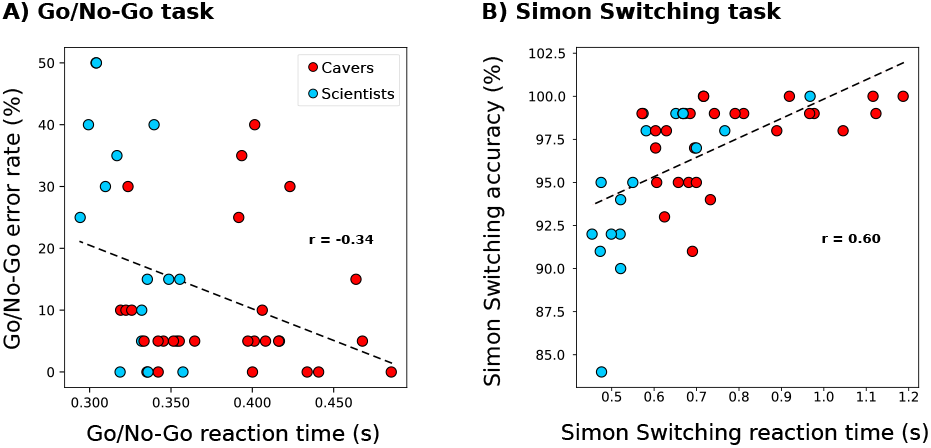
Reaction time on correct trials were A) negatively-related to error rates on the Go/No-Go task and B) positively-related to accuracy rates on the Simon task. Results show that error rate was low on average though varied considerably on Go/No-Go (cognitive inhibition) task, and accuracy rate was overall high on the Simon task, demonstrating that participants were able to do the tasks. r-values are included for descriptive purposes (data points are not independent).

**Figure 6:**
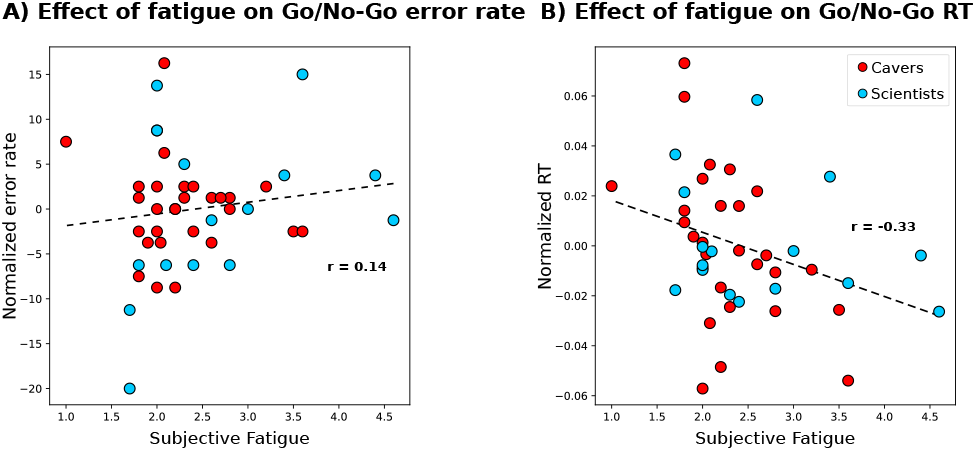
Effect of fatigue on both the Go/No-Go A) error rate and B) reaction time. In A), a trend is observed between decreased Go/No-Go accuracy (higher error rate) and decreased feelings of energy. In B), reaction times generally decreased in the Go/No-Go task when people reported greater fatigue. r-values are included for descriptive purposes (data points are not independent).

Using the performance results from the Simon Switching task, we can distinguish between two different cognitive costs (see Methods) and measure their impact on accuracy and reaction time: the ‘Simon effect’ (cost of stimuli position/direction incongruence) and the ‘Switch effect’ (cost of switching between instructions). On these metrics, lower negative numbers for accuracy or higher positive numbers for reaction time indicate worse performance on difficult trials (incongruent or switching trials, respectively). Our results suggest that increased fatigue may be related to decreased ‘selective attention’ when responding to incongruent stimuli. As shown in Figure 7A (left panel), a higher level of participant fatigue shows a trend towards better accuracy on congruent than incongruent trials (negative value of Simon effect). Similar trends were found when looking at the relationship between the Switch effect with subjective fatigue ratings, suggesting a similar decrease in cognitive flexibility with increased fatigue levels (see Figure 7B, left panel). In terms of reaction time, we do not see any trend of correlation with the effect of reaction time being close to 0 (meaning similar reaction time for both non-switching and switching trials (as seen in Figure 7A right panel) and both incongruent and congruent trials (as seen in Figure 7B, right panel)).

**Figure 7:**
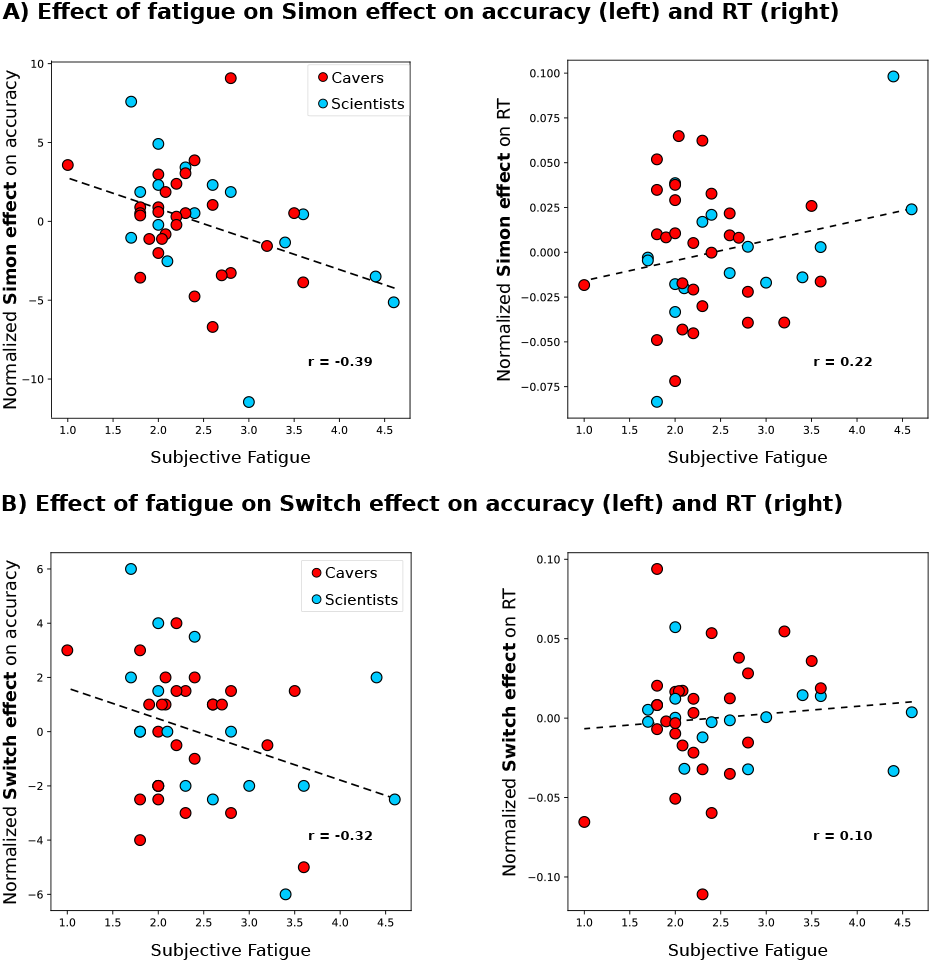
Effect of fatigue on both the A) switch effect and B)incongruence (Simon) effect on accuracy and reaction time for correct trials in the Simon Switching Task as a function of subjective fatigue. A similar trend of increased errors is observed for both the effect of switching and incongruence while fatigued. Note: This correlation plot serves as a visual representation and does not imply statistical significance. The provided data highlights trends and serves as an illustrative example of what could be explored with a larger dataset.

### 3.3. Subterranean measurements

#### 3.3.1. Video recordings

Team members did not find that wearing the body cameras interfered with activities, found them easy to turn off and on for privacy, and did not ask that any data be deleted. Aside from scratches, no equipment was damaged during the expedition, suggesting an adequate degree of robustness. After inspection of the data captured we conclude that the audiovisual recordings are of good quality and speech was clearly audible enabling use for further analysis. In some cases, cameras were not optimally-positioned to capture activity in front of the wearer, were occluded by clothing, or shaking or swinging movements reduced clarity. Due to the low-light conditions, infrared recordings were better than visual spectrum recordings (see Figure 8A for examples of still frames of subterranean footage).

**Figure 8:**
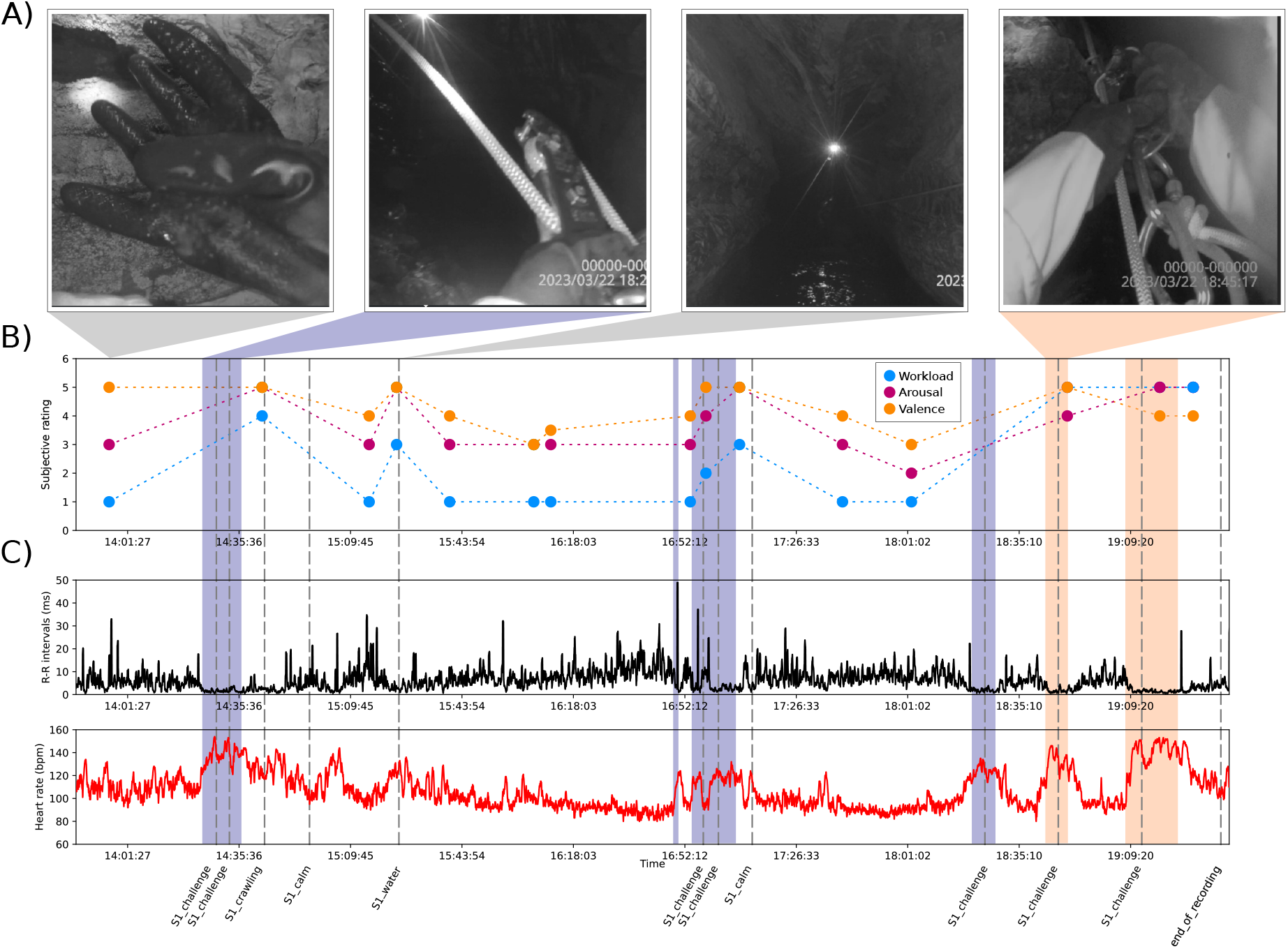
Example analysis from data recorded during cave progression. A) Audiovisual recordings captured using a body camera help researchers contextualize recordings and mark events of interest. Left to right, the participant: gestures to indicate subjective ratings of state; manipulates equipment to descend a vertical obstacle; prepares to negotiate a subterranean lake; and ascends using a rope and gear. B) Subjective ratings of Workload, emotional Arousal and emotional Valence (‘WAV’) given periodically by the participant during progression were extracted from the audiovisual recording. C) Cardiac (heart rate and R-R interval) data was simultaneously measured with a sensor worn on a chest belt. Heart rate increases and R-R interval decreases occur as a result of physical exertion, but also as a results of cognitive state via the autonomic nervous system, e.g., in anticipation of the lake crossing. Background colours represent moments of higher physical activity (blue: descending, orange: ascending)

#### 3.3.2. Subjective ratings

Both oral and gesture WAV ratings were clearly recorded (see first still frame of Figure 8A) and participants did not find the request intrusive. This technique allowed us to collect subjective data within the cave without any additional equipment. These metrics inform us about the perceived emotional and mental workload experienced by individual participants. For visualization purposes, we plotted the evolution of WAV ratings during a 6-hour exploration activity for one participant (see Figure 8B). We see interesting relationships between specific events (e.g., descending on the rope) and their impact on subjective ratings. It is likely necessary for members to be reminded to make ratings, possibly through regular notifications (experience sampling method; Csikszentmihalyi et al., 2014).

#### 3.3.3. Heart rate variability measures

Heart rate monitors were worn across the chest of participants wearing the body cameras and data was successfully aligned across devices. The method of aligning the electrocardiological data with the videos using the camera clocks was adequate for the current purpose, though synchronized data collection would be preferable. Figure 8 shows an example recording with key events noted. As suggested by visual inspection of the subjective ratings, we observe interesting relationships between change in cardiac activity as observed with R-R intervals and heart rate (see Figure 8C) and events of interest.

By evaluating the physical and cognitive workload of several participants’ activity using the footage from the body cameras, we were able to find periods of interest and observe trends of differences in HRV metrics between these periods (see Figure 9). Focusing on the stationary periods we observe trends in the data measured during this expedition, showing a decrease in the mean SDNN and mean pNN50 and an increase in LF/HF between the Calm and Anticipation conditions (as seen in Figure 9). The difference between the “Low” and “High” progression periods highlights the variability in the effect of physical activity on cardiac activity and HRV metrics. However, if we focus on the last two conditions where physical activity was similarly high but cognitive activity differed, we once again observe the decreasing trend in SDNN and pNN50, and an increase in LF/HF.

**Figure 9:**
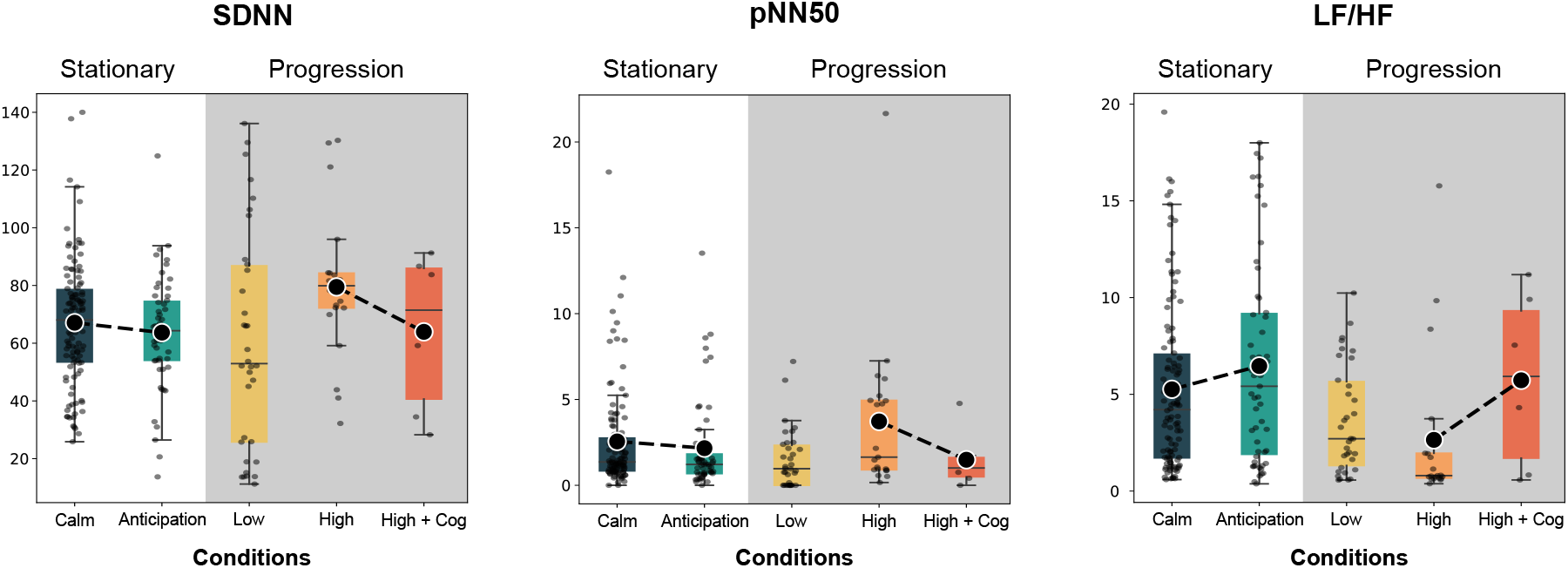
Heart rate variability metrics for stationary and cave progression periods with different levels of cognitive and physical activity. While stationary (white background), there were periods of low cognitive activity (Calm) and high cognitive activity (Anticipation). During cave progression (grey background), there were periods of low physical activity (Low), high physical activity (High), and high physical activity combined with high cognitive activity (High + Cog). The results suggest separability of user state, given knowledge of activity level.

### 3.4. Occasional measurements

#### 3.4.1. Sleep

We present summary physiological measures of sleep quality as indexed by EEG, across all nights recorded following exploration days in Supplementary Table 2. In general, recording quality was good despite the difficult conditions, establishing feasibility. Participants had more disturbed sleep as indexed by wake after sleep onset (WASO; mean: 38.0 min, SD: 29.6 min, minimum: 14 min, maximum: 101 min) and a higher number of awakenings than what would be expected for healthy participants sleeping in their home environments (mean: 23.9, SD: 4.3, minimum: 17, maximum: 30). The effect of the expedition environment is illustrated in Figure 10, which shows the sleep architecture (hypnograms) of one participant during a night recorded before the expedition and one during the expedition.

**Figure 10:**
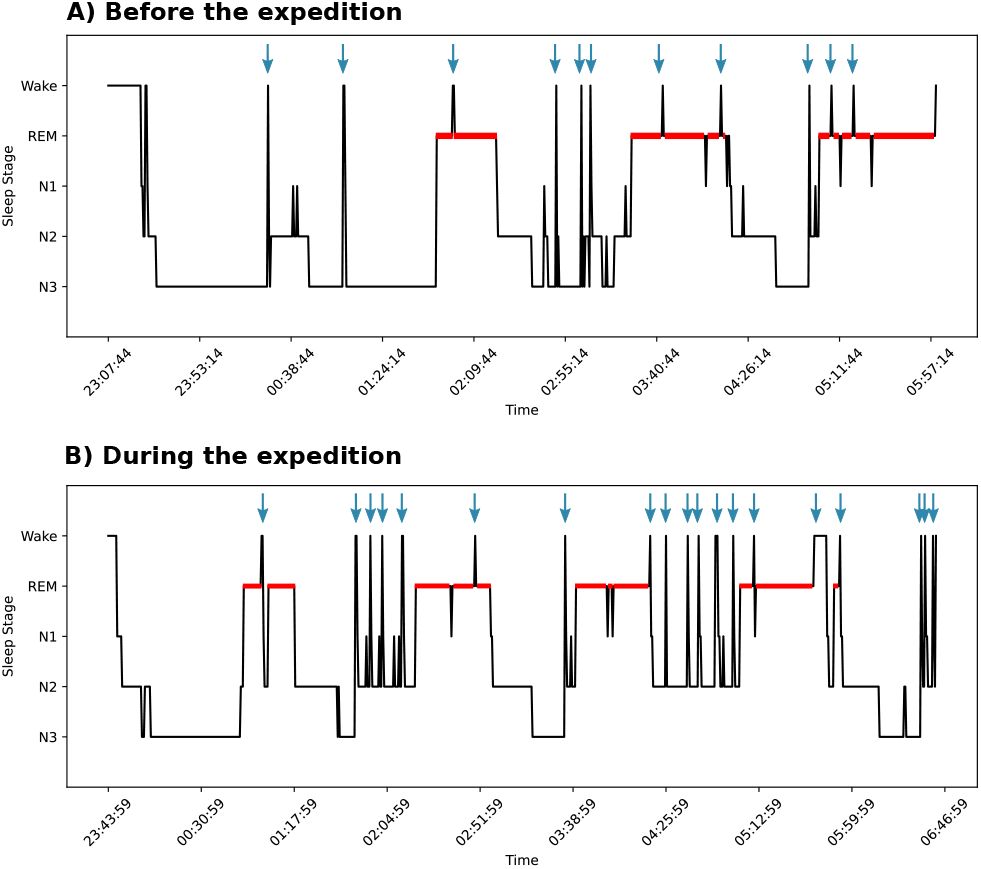
Impact of expedition environment on sleep structure. Example of a hypnogram for the same individual A) before the expedition and B) during the expedition. Red bars represent the duration of REM sleep and blue arrows indicates awakenings during the night, which are more numerous in B) and correspond to less deep and restful sleep. REM: rapid eye movement sleep; N1: stage 1 non-rapid eye movement (NREM) sleep (i.e., light sleep); N2: stage 2 NREM sleep; N3: stage 3 NREM sleep (i.e., deep sleep).

#### 3.4.2. Interviews

We recorded one general interview per participant, mostly on the last day (Figure 3); interviews took between 9 and 45 minutes. The portion of the interview in which the researchers go through the video with the participants proved impractical, as it was time-consuming and groups of speleologists often arrived to camp late, tired, and hungry, and had to cook, clean up, take care of equipment, as did the researchers. It was also impractical to complete the second part of the interviews, in which participants are asked about the day’s activities, each day given time limitations and only two interviewers. We had intended that this section provide additional information for interpreting and analyzing each day’s video recordings; however, a more efficient solution would be to have a brief written or audio log kept by each participant.

The interviews were most valuable in allowing us to understand some more nuanced aspects of speleology that could help contextualize our observations. For example, though some participants mentioned that caving was not a competition per se, many still expressed fear of being the slowest member of the team, and mentioned that certain aspects of caving could become competitive for certain people (e.g., comparing depths explored, speed of progression, quantity of discoveries, or personal performance). Still, participants tended to emphasize the importance of team cohesion, recognizing limits, and supporting each other, and emphasized the need to minimize competition and promote collaboration. Interviews also yielded insights into the different levels of expertise and experience in speleology, and individuals’ awareness of their own skills with respect to those of team members. For additional information from the interviews including general trends of responses among most participants, see Supplementary Materials.

## 4. Discussion

The main goal of this project was to test the feasibility conducting cognitive neuroscience and neuropsychology research in extreme environments, in this case during a speleological expedition. In general, the majority of the data collection techniques we used proved workable, with some caveats and adaptations to procedures necessary in future work. We summarize the findings regarding feasibility, adaptations, and recommendations in Table 1, and discuss them further in the next section.

### 4.1. Feasibility considerations and recommendations

#### 4.1.1. Questionnaires

The Emotional Response Questionnaire results showed a end towards participants having higher scores on cognitive eappraisal and lower scores on expressive suppression than community samples. Emotional regulation strategies are essential to ensure group cohesion and functionality on safety critical missions (A. Griffith et al., 2014). Inadequate emotional regulation may jeopardize missions through worsening performance when operating machinery (Hancock et al., 2012) or completing physically-demanding tasks (Lazarus, 2000), and degrading motor behaviour and attentional control (Beatty and Janelle, 2020). For these reasons we may wish to study differences in emotional regulation strategies, as well as for their effects on physiological measures, group dynamics, and mission success. Although they have previously been characterized as ‘adaptive’ or ‘maladaptive’, different strategies may be more or less effective depending on the expedition context and the goals of the emotional regulation (Wagstaff, 2014; Wagstaff and Weston, 2014; for example, the use of some strategies may facilitate group cohesion, while others could be better suited for individual goals in high-risk situations).

A drawback to the ERQ is that it is unclear whether self-reported personal tendencies in emotion regulation strategies were actually used during the expedition, which presents different challenges to those experienced in daily life. Future directions for exploring emotional regulation in cavers might include asking them to reflect on specific use of strategies during expeditions, and comparing cavers’ skills and traits with other groups (i.e., astronauts, general population), which could be done in larger groups outside of the expedition setting (i.e., societies or organizations’ members).

The “Psychological Skills Inventory for Caves” questionnaire was adapted from a similar tool in sports psychology, and we aimed to assess its applicability. Our participants were encouraged to provide feedback on the items in the questionnaire for future refinement. For example, a few members indicated that the items related to mental imagery did not apply to them; however, others scored quite high (up to 95%) on the mental rehearsal subscale. These items were therefore left in the questionnaire (with slight reformulations based on feedback from other participants). Overall, the modifications made to the questionnaire post-feedback demonstrate the iterative nature of questionnaire development and the importance of incorporating the perspectives of the target population. Wording modifications may affect the underlying constructs being measured, and further factor analyses are warranted to examine whether the PSIC retains the same factor structure as the original Psychological Skills Inventory for Sports (PSIS). The original (pre-expedition) and updated (post-feedback) lists of questions in the PSIC are available as Supplementary Materials, in both French and English.

#### 4.1.2. Cognitive tasks

The Go/No-Go and Simon Switching tasks proved feasible in terms of their timing and equipment demands. Despite the sub-optimal testing conditions (crowding, distraction), both cognitive tasks showed trends suggesting they were sensitive to subjective experiences of fatigue (Figures 6 and 7). Results on the selected cognitive tasks did not exhibit a ceiling effect or show much evidence of learning, suggesting that both tasks are suitable for repeated, longer-duration use.

We noted some interesting discrepancies with previous work as regards the relationship between fatigue levels and reaction time. In laboratory-based work, reaction times are generally similar or slower when participants are fatigued (e.g., Demos et al., 2016; Lee et al., 2022); our results instead suggest that participants respond faster yet accept lower accuracy scores when tired (Wickelgren, 1977). This finding may have implications for safety, should team members make a similar trade-off during movement, e.g., while following procedures to ascend from a long day of exploration. Although cavers are likely to prioritize accuracy in safety-relevant tasks, statistics from aviation suggest that accidents are most frequent during approach and landing phases, during which high task load coincides with fatigue (Kharoufah et al., 2018). Previous work suggests that even highly-trained professionals with a clear understanding of safety consequences are affected by such cognitive changes. These results hint at a benefit to studying cognition in non-laboratory conditions; relationships that do affect real-world outcomes may not arise under ideal laboratory conditions.

We can conclude that these tasks and similar computerized cognitive tasks that measure other aspects of cognition (e.g., sustained attention) are feasible on expeditions, if kept short. Given small sample sizes and inter-individual variability in performance and in the speed-accuracy trade-off, the most powerful statistical designs would necessitate as many testing points as possible (e.g. daily for the entire expedition), and perhaps a pre-expedition and post-expedition baseline period (which could be run remotely using an online platform, in the case of a geographically-distributed team). As regards practical applications, we believe that cognitive tests are unlikely to be directly useful to determine individuals’ daily state for team decision-making purposes; subjective fatigue reports do correlate with performance and require less time and effort.

#### 4.1.3. Bodycams

We determined the use of bodycams as feasible for cave exploration: the participants were comfortable with the equipment, the video (infrared) and audio quality were sufficient, and the cameras were sufficiently sturdy for the activity. The videos allowed researchers to identify events of interest and to collect data points using the WAV system; however, this process was facilitated by notes from interviews and first-hand knowledge of the expedition. It was difficult for off-site team members to interpret the videos independently, supporting the need for embedding scientists within expeditions, or asking participants to record activities in written or oral logs.

#### 4.1.4. Heart rate variability

Collecting usable heart rate data was feasible during the cave expedition. The equipment fit well under other clothing and did not hinder movement. We were able to align heart rate data with the body camera recordings, although collecting precisely synchronized data would be preferable. Combining heart rate data along with subjective measures such as WAV ratings allowed us to get a more complete understanding of the human experience of cavers, as we were able to use those ratings to pinpoint events of interest. Cardiac measures are also influenced by movement, and dissociating the effects of workload or emotional state from those induced by physical activity is an unresolved challenge of high interest for researchers working on measuring mental states in real-world environments (e.g., Albuquerque et al., 2020). It is worth noting that mental and physical effects on cardiac measures may be tightly correlated *because* movement requires cognition. Studies have shown that even merely walking uses higher order cognitive and cortical control mechanisms (Mirelman et al., 2014), suggesting that moving within a cave environment would be correlated with cognitive resource use and increased mental workload.

#### 4.1.5. Subjective ratings

The subjective fatigue scale took very little time and was practical to administer, noting that care should be taken to avoid team interactions that might affect people’s ratings (e.g., favour written or computerized format as opposed to oral ratings). As fatigue scores seem to correlate somewhat with cognitive performance, daily ratings may be useful to help individuals, team leaders, or the entire group make safe decisions about exploration activities. However, it is important to note that there can be discrepancies between self-evaluations of mental state and actual cognitive performance (Berastegui et al., 2020; Boardman et al., 2021), particularly as regards people’s evaluations of their own capability for performance when poorly rested, suggesting that scientific investigations should include both subjective and objective measures.

#### 4.1.6. Sleep

It was feasible to objectively record sleep duration and quality in the expedition environment. Because the headbands measure EEG, the data can be used to extract additional fine-grained information like number of sleep spindles and slow oscillations (Debellemaniere et al., 2018). Limitations to the specific tool used (Dreem) are that it has limited data storage capacity (several nights) and internet connectivity is required to remove and save the data, and that its cloth exterior is difficult to clean in camp conditions. For research questions that do not require fine-grained sleep measures, a sports watch that can approximate sleep duration through accelerometers and cardiac measures may be sufficient to quantify rest of expedition members (Miller et al., 2022). In future work, sleep metrics can be used to explore relationships to next-day fatigue level and cognitive performance, and could be used as input to expedition decisions. As with the cognitive tasks, small sample size limitations can be overcome with repeated measures per member, a pre- and post-expedition baseline period, and statistical approaches that take into account within- and between-subjects variability.

#### 4.1.7. Interviews

Qualitative interview data provided valuable leads for data analysis and interpretation. Instances of increased cognitive labour or emotional arousal were described and reported, indicating relevant data points in the cardiac data and providing greater descriptive depth to emotional regulation and cognitive workload measures. Participants were able to recount challenging moments of a day’s caving activities, and explain phenomenological aspects of the experience, providing information that may have otherwise been missed. Furthermore, subjective perceptions of difficulty of different aspects of caving varied by participant, and interviews provided insights into hidden social, emotional, and cognitive experiences (fear of slowing the group down, perception of team dynamics, etc.).

Conducting interviews during expeditions has unique challenges due to the limited time and personnel available, logistical demands, and participants’ (and researchers) physical and mental fatigue during periods at the camp. We suggest separating questions on general background and motivation that can be answered before or after the expedition (remotely if necessary), standardizing interview questions, and providing interviewers with training on techniques. Written or oral daily logs could also be set up for independent use by participants (e.g., using the body cameras’ audio recording function), to reduce the bottleneck of interviewer availability and improve flexibility of scheduling during camp periods. In-person interviews could then concentrate on exploring topics of interest related to specific exploration events.

## 5. Overcoming measurement limitations inherent to expeditions

The small size heterogeneity of the team in the present work is likely representative of future expeditions, meaning that despite refining our tools and measures, major challenges to conducting empirical work in expedition environments remain. We posit that these obstacles can be partly overcome by measuring multiple data-points within-participants (i.e., measuring cognitive tasks for the entire expedition as well as pre- and post-expedition baselines); perhaps using statistical models that consider both within- and between-participants’ sources of variance (e.g., linear mixed effects models; Pinheiro and Bates, 2006); and by deliberately combining measures and approaches to challenge or confirm findings, a technique sometimes called ‘triangulation’ (Turner and Turner, 2009).

The data collection methods evaluated in this work can be categorized into quantitative approaches, including the questionnaires, scales, and cognitive tasks; and qualitative approaches, in the form of interviews and behavioural observations that were made by reviewing the audiovisual recordings. Although not a commonly employed technique in cognitive neuroscience research, these ‘mixed methods’ approaches are used in psychological research and are being increasingly applied in other areas like healthcare (Dures et al., 2011; Shorten and Smith, 2017) and education (Clark, 2019). A key concept in mixed-methods research is that information from both approaches is integrated and linked to enable a more panoramic view of the ‘research landscape’ (Shorten and Smith, 2017). We implicitly used a mixed-methods approach, for example by using interviews and direct observation to help annotate the videos and analyze the physiological data; however, an explicit application of mixed-methods principles and best practices could help field researchers manage complexity, small sample sizes, and difficulty replicating exact conditions, which make purely quantitative approaches less robust. Researchers considering using mixed-methods approaches may wish to familiarize themselves with some of the theoretical and practical challenges that come with mixing research approaches (Bishop, 2015; Dures et al., 2011), as well as best practices for specifying an appropriate mixed methods design (Clark, 2019; Creswell and Clark, 2017; Hong et al., 2018).

## 6. Conclusions

The results of this first cognitive science and neuropsychology study in real expedition conditions supports feasibility of future work, and takes several practical steps towards more robust data collection in the field, with implications for other safety-critical environments such as remote planetary exploration and search-and-rescue operations (Bartlett and Cooke, 2015; Hart et al., 2022; St-Onge et al., 2020, 2019). More broadly, we hope to contribute to a new movement in which studying complex, challenging, naturalistic behaviours in humans complements laboratory work to contribute to our understanding of brain function and behaviour (Nastase et al., 2020; Shemesh and Chen, 2023; Smith, 2023).

## Data availability statement

Anonymized data for the questionnaires and cognitive tasks are available on Open Science Foundation (https://osf.io/a9q6f/?view_only=e7cf482d34784cf58908f996ff7b856a), as are copies of the software for the cognitive tasks, questionnaires, and semi-structured interview questions. Videos and interviews are not available as they cannot be anonymized.

## CRediT author statement

Anita Paas: Conceptualization, Methodology, Formal analysis, Writing - Original Draft, Writing - Review & Editing, Visualization. Hugo Jourde: Conceptualization, Methodology, Data collection, Formal analysis, Data Curation, Writing - Original Draft, Writing - Review & Editing, Visualization. Arnaud Brignol: Conceptualization, Methodology, Formal analysis, Writing - Original Draft, Writing - Review & Editing, Visualization. Marie-Anick Savard: Conceptualization, Methodology, Formal analysis, Data Curation, Writing - Original Draft, Writing - Review & Editing, Visualization. Zseyvfin Eyqvelle: Formal analysis, Writing - Original Draft, Writing - Review & Editing, Visualization. Samuel Basseto: Conceptualization, Writing - Review & Editing, Project administration, Funding acquisition. Giovanni Beltrame: Conceptualization, Writing - Review & Editing, Project administration, Funding acquisition. Emily Coffey: Conceptualization, Methodology, Data collection, Formal analysis, Data Curation, Writing - Original Draft, Writing - Review & Editing, Visualization, Supervision, Project administration, Funding acquisition.

## Acknowledgements

The authors thank Mexpé team members and organizers, and Spéléo Québec, for inviting us to participate in Mexpé 2023, providing technical training, equipment, organization and logistics, and maintaining a safe and secure expedition environment, and for participating as collaborators and participants in this research. The authors would also like to thank Daniel Caron for facilitating access to the Saint-Léonard Cave in Québec, Canada, for pre-expedition equipment testing; Keelin Greenlaw for assisting with the semi-structured interview questions; Koresh Khateri for technical support; and the committee members and staff of the Concordia University Human Research Ethics Committee for constructive feedback and timely approval. EC and GB are supported in part by funding from the Government of Canada’s New Frontiers in Research Fund (NFRF). We also acknowledge the support of the Canadian Space Agency (CSA) [21FACONB14]. ZE received a Research for Undergraduates in Space Health (RUSH) Award from the Canadian Space Health Research Network.

## Footnotes

1 https://osf.io/a9q6f/?view_only=e7cf482d34784cf58908f996ff7b856a

2 https://gitlab.pavlovia.org/vuandre1/goNogoAlpha4

3 https://gitlab.pavlovia.org/xiangjie/simon-switching-task-1

4 https://dreem.com/

## Notes

### Competing Interest Statement

The authors have declared no competing interest.

